# Genomic characterization of novel bat kobuviruses in Madagascar: implications for viral evolution and zoonotic risk

**DOI:** 10.1101/2024.12.24.630179

**Authors:** Freddy L. Gonzalez, Hafaliana Christian Ranaivoson, Angelo Andrianiaina, Santino Andry, Vololoniaina Raharinosy, Tsiry Hasina Randriambolamanantsoa, Vincent Lacoste, Philippe Dussart, Jean-Michel Héraud, Cara E. Brook

## Abstract

Kobuviruses (family *Picornaviridae*, genus *Kobuvirus*) are enteric viruses that infect a wide range of both human and animal hosts. Much of the evolutionary history of kobuviruses remains elusive, largely due to limited screening in wildlife. Bats have been implicated as major sources of virulent zoonoses, including coronaviruses, henipaviruses, and filoviruses, though much of the bat virome still remains uncharacterized. While most bat virus research has historically focused on immediately recognizable zoonotic clades (e.g. SARS-related coronaviruses), a handful of prior reports catalog kobuvirus infection in bats and posit the role of bats as potential progenitors of downstream kobuvirus evolution. As part of a multi-year study, we carried out metagenomic Next Generation Sequencing (mNGS) on fecal samples obtained from endemic, wild-caught Madagascar fruit bats to characterize potentially zoonotic viruses circulating within these populations. The wild bats of Madagascar represent diverse Asian and African phylogeographic histories, presenting a unique opportunity for viruses from disparate origins to mix, posing significant public health threats. Here, we report detection of kobuvirus RNA in Malagasy fruit bat (*Eidolon dupreanum*) feces and undertake phylogenetic characterization of one full genome kobuvirus sequence, which nests within the Aichivirus A clade - a kobuvirus clade known to infect a wide range of hosts including humans, rodents, canids, felids, birds, and bats. Given the propensity of kobuviruses for recombination and cross-species infection, further characterization of this clade is critical to accurate evaluation of future zoonotic threats.

## Background

Picornaviruses in the viral family *Picornaviridae* are non-enveloped RNA viruses that infect a wide range of vertebrates, from birds and fish to a variety of mammals, including both humans and bats^1,2^. Famed human picornaviruses include poliovirus (genus: *Enterovirus*), which causes the paralyzing human disease poliomyelitis^3^, and the rhinoviruses (also genus: *Enterovirus*) which cause the common cold^4,5^. Arguably the most well-known animal picornavirus is the first described in this clade, Foot-and-Mouth-Disease Virus (genus: *Apthovirus*), which causes the agriculturally-devastating disease of the same name in cloven-hoofed animals^6^. To date, bat picornaviruses have been generally overlooked as potential zoonotic pathogens due to the lack of documented zoonotic spillover events in this clade^7^. This oversight has resulted in limited bat picornavirus surveillance which hinders efforts to describe their evolutionary history. In contrast, bat coronaviruses have garnered significant attention due to their established zoonotic potential. Unlike coronaviruses, which exhibit tight coevolutionary signatures with specific bat host species, picornaviruses show low host specificity, as highly similar variants have been detected across a wide array of bat species, suggesting a more generalized host range^2,8^. Further screening for bat picornaviruses in high-risk areas of wildlife-human interaction will provide crucial insights into their evolutionary history and potential for cross-species transmission.

Kobuviruses represent one clade of many recently discovered enteric picornaviruses known to cause severe gastroenteritis in humans and animals^7^. As a clade, they are subdivided into genotypes Aichivirus A-F. Genotypes falling under the Aichivirus A classification are hosted by humans, canids, rodents, felids, birds, and bats^9–11^, and those within the Aichivirus B-F classification are hosted by cattle, swine, sheep, rabbits, and bats^12–17,11,18^. Structurally, kobuviruses are small (∼30-32 nm), icosahedral, non-enveloped viruses with a single-stranded positive sense RNA genome of 8.2-8.4 kb in length^18^. They contain only one open reading frame (ORF), which encodes three structural proteins (VP0, VP3, and VP1) and eight nonstructural proteins (L, 2A, 2B, 2C, 3A, 3B, 3C, and 3D)^19^. These genomic and structural features not only play a critical role in the ability of kobuviruses to infect a wide range of hosts but also provide valuable insight into their evolutionary adaptability and potential for zoonotic transmission.

Madagascar is home to 51 species of bat^20^, many of which are endemic and have undergone long evolutionary divergence from sister species in both Africa and Asia^21,22^. Recent evidence identifies Madagascar fruit bats as hosts for numerous circulating viruses^23–27^, some of which are potentially zoonotic. Additionally, longitudinal serological surveillance shows that female Malagasy bats exhibit elevated antibody titers during the periods of gestation and reproduction, suggesting that their exposure to viruses such as henipaviruses and filoviruses may be seasonally linked. Routine bat virus surveillance is thus a critical public health priority, particularly in regions with high human-bat contact rates^28^. With rapidly changing ecology, urbanization, climate change, increased travel, and fragile public health systems, the frequency of bat zoonoses is likely to rise^29^. Despite growing evidence of viral circulation in Madagascar’s diverse bat populations, the prevalence and evolutionary history of many viral taxa—including kobuviruses—remain largely unexplored in these hosts.

Here, we carried out metagenomic Next Generation Sequencing (mNGS) of RNA extracted from fecal samples collected from three endemic Malagasy fruit bat species (*Pteropus rufus, Eidolon dupreanum, Rousettus madagascariensis*). We present the first detection and characterization of any kobuvirus circulating within Malagasy fruit bats and use phylogenetic tools to demonstrate that Malagasy fruit bat kobuviruses share immediate ancestry with previously described members of kobuvirus sub-clade Aichivirus A, a clade that includes several human-infecting species. We aim for our descriptive work to provide a baseline on which future work will build understanding of kobuvirus dynamics within Malagasy wildlife hosts and evaluate their potential capacity for cross-species transmission.

## Materials and Methods

### Bat Sampling

As part of a multi-year study examining the dynamics of potentially zoonotic viruses in three endemic species of Madagascar fruit bats (*Pteropus rufus, Eidolon dupreanum, Rousettus madagascariensis*), bats were captured monthly in species-specific roost sites in the Districts of Moramanga and Manjakandriana, Madagascar between 2018 and 2019 (*P. rufus*: Ambakoana roost, −18.513 S, 48.167 E; *E. dupreanum*: Angavobe cave, −18.944 S, 47.949 E; Angavokely cave = −18.933 S, 47.758 E; *R. madagascariensis*: Maromizaha cave, −18.9623 S, 48.4525 E). Bats were live-captured using nets hung in tree canopies (*P. rufus*) and over cave mouths (*E. dupreanum, R. madagascariensis*) at dusk (17:00-22:00) and dawn (03:00-07:00). Captured bats were manually restrained, and bat sex, species, and age (juvenile or adult) were morphometrically determined in the field following previously published protocols^23,30–32^. Fecal swabs were collected from all captured individuals, placed into viral transport medium, and frozen in liquid nitrogen. After sampling, swabs were transported to −80^0^C freezers at the Virology Unit at Institut Pasteur de Madagascar for long-term storage. In total, 690 bats were captured (*P. rufus*: 68, *E. dupreanum*: 288, *R. madagascariensis*: 334).

This study was carried out in strict accordance with research permits obtained from the Madagascar Ministry of Forest and the Environment (permit numbers 019/18, 170/18, 007/19) and under guidelines posted by the American Veterinary Medical Association. All field protocols employed were pre-approved by the UC Berkeley Animal Care and Use Committee (ACUC Protocol #AUP-2017-10-10393), and every effort was made to minimize discomfort to animals.

### RNA Extraction

A random subset of fecal samples distributed across all three species was selected for downstream molecular analysis, including RNA extraction and mNGS (*P. rufus:* 26 male/18 female*, E. dupreanum:* 52 male/93 female*, R. madagascariensis:* 49 male/47 female) (**Table 1**). Samples undergoing mNGS corresponded to captures in Feb-Apr, Jul-Sep, and December 2018 or in January 2019. RNA was extracted at the Virology Unit at the Institut Pasteur de Madagascar, using the Zymo Quick DNA/RNA Microprep Plus kit (Zymo Research, Irvine, CA, USA), adhering to the manufacturer’s instructions while also including a DNAse digestion step. Water controls were extracted in conjunction with samples on each extraction day. Post-extraction, RNA underwent quality control on a nanodrop to assess its purity. A 260/280 ratio absorbance that did not exceed 2 was used to ensure that a quantifiable concentration was present. Extractions that passed screening were stored in freezers at −80^0^C and transported to the Chan Zuckerberg Biohub (San Francisco, CA, USA) for library preparation and mNGS.

**Table 1.** Positive Kobuvirus Samples. Summary table showing total bats captured by species and location from a random subset of fecal samples subject to mNGS.

| Roost site | Species | Total<br>sampled by<br>site (n =<br>285) | Total<br>Kobuvirus<br>positive (n =<br>2) | Total<br>sampled<br>(Male,<br>Female) | Total<br>Kobuvirus<br>positive<br>(Male,<br>Female) |
| --- | --- | --- | --- | --- | --- |
| Ambakoana | *Pteropus rufus* | 37 | 0 (0%) | 23,14 | 0,0 (0%, 0%) |
| Angavobe | *Eidolon*<br>*dupreanum* | 37 | 1 (2.7%) | 11,26 | 0,1 (0%,<br>3.84%) |
| Angavokely | *Eidolon*<br>*dupreanum* | 108 | 1 (0.93%) | 41,67 | 0,1 (0%,<br>1.49%) |
| Maromizaha | *Rousettus*<br>*madagascariensis* | 96 | 0 (0%) | 49,47 | 0,0 (0%, 0%) |
| Mahialambo | *Pteropus rufus* | 7 | 0 (0%) | 3,4 | 0,0 (0%, 0%) |

### Library Preparation and mNGS

Four randomly selected samples from each bat species (*Pteropus rufus, Eidolon dupreanum, Rousettus madagascariensis*) underwent additional quantification using an Invitrogen Qubit 3.0 Fluorometer and the Qubit RNA HS Assay Kit (ThermoFisher Scientific, Carlsbad, CA, USA). Following quantification, RNA samples, and water samples from prior extraction, were pipetted into 96-well plates to automate high throughput mNGS library preparation. Based on initial quantification, 2uL aliquots from each plated sample were diluted 1:9 on a Bravo liquid handling platform (Agilent, Santa Clara, CA, USA). 5uL aliquots from each diluted sample were pipetted into 384-well plates for mNGS library preparation. Fecal samples were arrayed on distinct 384 well plates for sequencing runs. Each 384-well plate included additional RNA samples isolated from cultured HeLa cells and lab water samples, which served as controls for library preparation. Samples were transferred into a GeneVac EV-2 (SP Industries, Warminster, PA, USA) to evaporate to conduct miniaturized mNGS library preparation with the NEBNext Ultra II RNA Library Prep Kit (New England BioLabs, Beverly, MA, USA). Library preparation was performed following manufacturer’s instructions with a few modifications: 25pg of External RNA Controls Consortium Spike-in mix (ERCCS, Thermo-Fisher) was added to each sample prior to RNA fragmentation, input RNA mixture was fragmented for 8 minutes at 94^0^C prior to reverse transcription, and a total of 14 cycles of PCR with dual-indexed TruSeq adapters was applied to amplify the resulting individual libraries. Resulting library pools then underwent quality and quantity measurements via electrophoresis (High-Sensitivity DNA Kit and Agilent Bioanalyzer; Agilent Technologies, Santa Clara, CA, USA), real-time quantitative polymerase chain reaction (qPCR) (KAPA Library Quantification Kit; Kapa Biosystems, Wilmington, MA, USA), and small-scale sequencing (2x146bp) on an iSeq platform (Illumina, San Diego, CA, US). Equimolar pooling of individual libraries from each plate was performed before running large-scale paired-end sequencing (2x146bp) on an Illumina NovaSeq sequencing system (Illumina, San Diego, CA, USA). The pipeline used to separate individual library outputs into FASTQ files of 146bp paired-end reads can be found at https://github.com/czbiohub-sf/utilities.

### Prevalence of Kobuvirus Sequence Detection in Field Specimens

Raw reads recovered from Illumina sequencing were host-filtered, quality-filtered, and assembled on CZID (v3.10, NR/NT 2019-12-01), an open-source, cloud-based *de novo* assembly pipeline for microbial mNGS data^33^, using publicly available full-length bat genomes from GenBank at the time of sequencing (July 2019) as the host background model. Samples were deemed kobuvirus positive if CZID assembled at least two contigs with an average read depth >2 reads/nt that showed significant nucleotide or protein BLAST alignments (alignment length >100 nt/aa and E-value < 0.00001 for nucleotide BLAST/ bit score >100 for protein BLAST) to kobuvirus reference sequences contained within NCBI NR/NT databases.

Offline BLASTn/x analyses of non-host contigs were conducted to cross-validate our search with CZID using a custom database of kobuvirus sequences from NCBI (last accessed: October 2021). Prior to BLAST searches, contigs were first deduplicated to remove redundant sequences using CD-HIT^34^ (v.4.8.1). BLAST hits from both searches identified two key contigs. One of these, a novel, full-genome length sequence that we eventually submitted to GenBank under accession number OP287812, was used as a reference for a third and final search aimed at filtering out low-quality hits from our prior runs.

### Genome quality assessment and annotation

We used CheckV^35^ (v1.0.1) to estimate genome completeness and potential host contamination of our putative genome (OP287812). We visualized OP287812 in Geneious Prime (V.2023.0.1) and aligned to previously annotated kobuvirus sequences obtained from NCBI with MAFFT^36^ (v.1.5.0). Protease cleavage sites were identified and used to define individual proteins, with NCBI sequences serving as references. We assumed that the 5’ and 3’ ends encompassed regions flanking the single open reading frame.

### Sequence similarity search

We conducted BLASTn and BLASTx searches to identify similarities between OP287812 and NCBI’s database of kobuvirus sequences within Geneious. We organized BLAST hits using Geneious Prime’s grade metric, a measure that produces a weighted score for hits composed of E-value, pairwise identity, and coverage to create a list of top 10 BLAST hits. We also generated an alignment between our full-(OP287812) and partial-length (OR082796) kobuvirus sequences, as well as with previously described bat kobuviruses, which we summarized using NCBI MSA Viewer (v.1.25.0).

### Sequence similarity analysis

We generated nucleotide and amino acid similarity plots comparing OP287812 to publicly available kobuvirus sequences recovered from NCBI (Accessions: KJ934637 and NC_001918). MAFFT sequence alignments were used as input for PySimPlot^37^ (v.0.1.1) using default window (Default: 100) and step sizes (Default: 1). Further data analyses and visualizations were carried out in RStudio^38^ (v.2024.04.2+764) using the tidyverse^39^ suite (v.2.0.0).

### Phylogenetic Analysis

We integrated our novel sequences with publicly available sequences on NCBI to perform three separate phylogenetic analyses: (a) a picornavirus maximum-likelihood (ML) tree spanning a conserved 7,000bp region, (b) a kobuvirus-only ML tree spanning a conserved 4,500bp region, and (c) a time-resolved Bayesian kobuvirus-only phylogeny spanning a conserved 5,500bp region. Sequences were aligned with MAFFT under default parameters and subjected to ModelTest-NG^40^ (v.0.1.7) to determine the best fit nucleotide substitution models to describe evolutionary relationships within each respective alignment. ML trees were built using RAxML^41^ (v.8.2.13) and visualized within RStudio using the ggtree^42^ (v3.16) package. Following standard practice outlined in the RAxML-NG manual, we computed 20 tree searches using 10 random and 10 parsimony-based starting trees under default heuristic search parameters for each original alignment, then selected the best-scoring topology. MRE-based bootstrapping tests were performed after every 50 replicates^43^, following Felsenstein’s method^44^, terminating at 1,000 bootstrap replicates. A similar approach was used to construct our Bayesian phylogenetic tree with BEAST2^45^ (v2.6.3), with the key difference being that representative sequences were selected from across the Kobuvirus-only ML phylogeny using Parnas^46^ (v.0.1.4) to ensure adequate coverage of tree diversity. More details for the generation of each phylogeny are available in our open-access GitHub repository (see Data Availability).

### Nucleotide Sequence Accession Number

We submitted both our annotated full-length genome sequence (8,263 bp) and partial-length sequence (2,077 bp) to NCBI where they were, respectively, assigned accession numbers: OP287812 and OR082796. Detailed descriptions of analyses are available on our GitHub (https://github.com/brooklabteam/Madagascar-Bat-Kobuvirus).

## Results

### Sequencing of fecal samples from Malagasy bats reveals kobuvirus prevalence and identification of a full-length genome

Two (2/285) fecal samples sequenced were kobuvirus positive via offline BLAST analyses (0.70% positivity) (**Table 1**), each originating from a different individual *Eidolon dupreanum* bat. Samples collected from *P. rufus* and *R. madagascariensis* did not demonstrate any evidence of kobuvirus infection.

The single full-length kobuvirus genome (OP287812) was identified in a sample collected from a juvenile *E. dupreanum* female in Angavokely cave, while the partial-length genome (OR082796) was identified in a sample collected from a non-lactating female adult *E. dupreanum* in Angavobe cave. Sequence OP287812 is 8,263 bp in length and was designated as ‘high quality’ by CheckV (completeness = 100, contamination = 0, CheckV quality = High quality, MIUVIG Quality = High quality). This sequence represents the most complete bat kobuvirus genome identified to date and the first bat kobuvirus to be identified in Madagascar.

### Genome annotation and comparative genomic analysis of the full-length kobuvirus genome OP287812 reveals homology across clades

We annotated the single ORF that spans OP287812 (7,305 nt and 2,435 aa) and identified protease cleavage sites across the genome (**Fig. 1A**). Reference sequences used to annotate the OP287812 genome can be found on **Supplemental Table 1**. Predicted cleavage sites occurring at junctions between L and VP0 (Glutamine/Glycine), 2A and 2B (Glutamine/Glycine), 2B and 2C (Glutamine/Glycine), 3A and 3B (Glutamine/Serine), and 3C and 3D (Glutamine/Serine) were consistent with prior findings in kobuviruses carried by other hosts^47,48^, suggesting considerable conservation of genomic content despite diverse host species.

**Figure 1.**
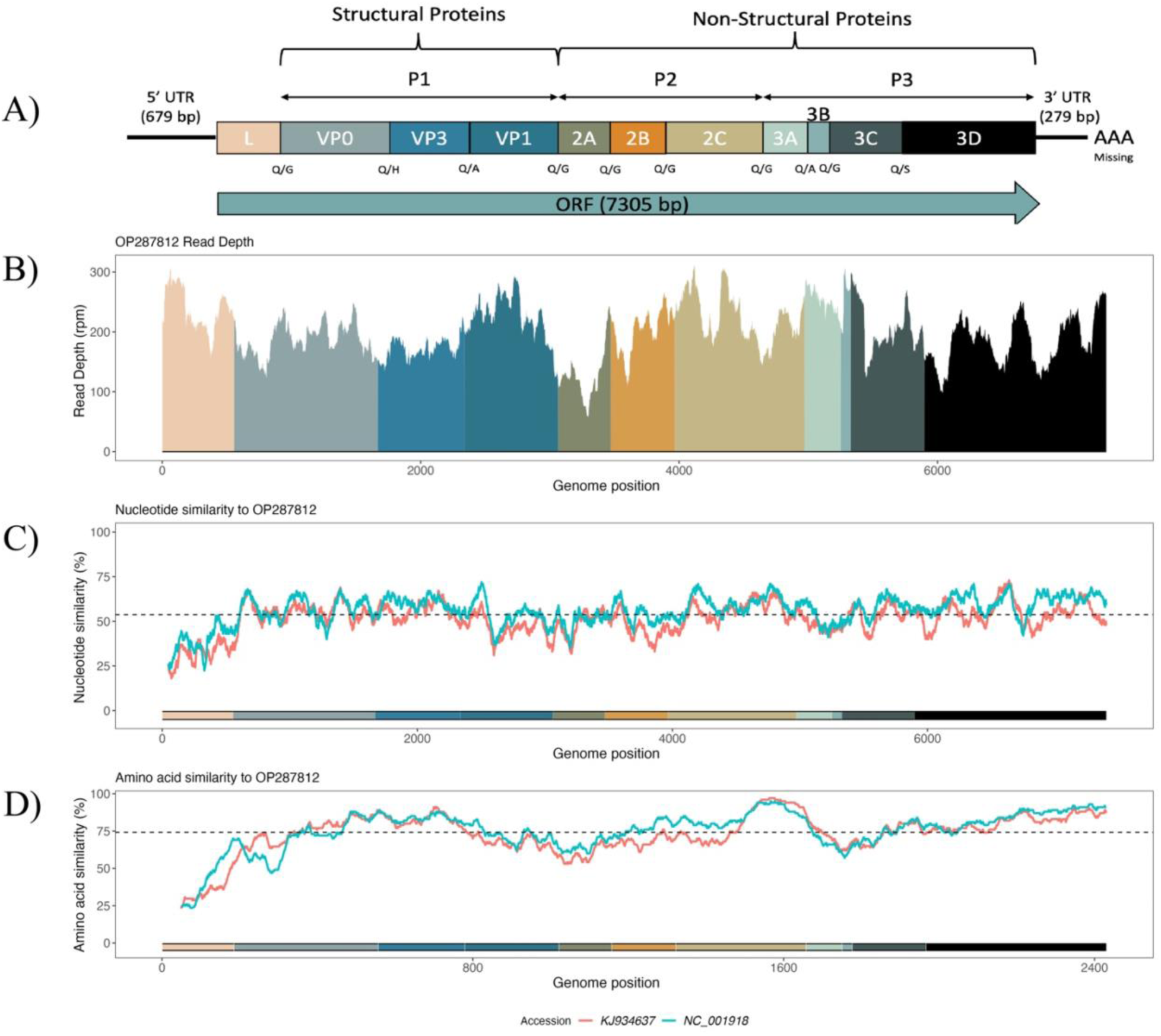
Genome Annotation and Similarity plot for OP287812. **A)** Genome annotation for OP287812 is shown at the top, with amino acid cleavage sites indicated below their corresponding proteins. Proteins are color-coded and displayed in panels **B-D**. **B)** Coverage map across OP287812’s open reading frame (ORF), excluding the 5’ and 3’ UTRs. Sequence depth is represented in reads per million (rpm), with colors indicating the proteins along the ORF. **C)** and **D)** Nucleotide and amino acid similarity plots, with OP287812 as the reference. The plots compare OP287812 to the most derived human kobuvirus sequence (NC_001918), and the most basal avian kobuvirus (KJ934637), in subsequent phylogenetic trees. Dashed lines indicate the average percent similarity between OP287812 and the query sequences.

We then plotted genome similarity between OP287812 and the most basal Aichivirus A variant, as well as between OP287812 and the most derived Aichivirus A variant included in our phylogenies **(Fig. 1C-1D**). We found that OP287812 shares an average nucleotide identity of 53.70% (**Fig. 1C**) and an average amino acid sequence similarity of 74.19% (**Fig. 1D**) to previously described kobuviruses (NC_001918 - Human and KJ934637 – Avian), consistent with BLAST results in which we recover high genome similarity to Aichivirus A variants.

We conducted additional BLAST searches of OP287812 against publicly available sequences in NCBI to assess its genomic similarity to previously identified kobuviruses (**Table 2**). Whole-genome BLASTn searches revealed a top hit to a human kobuvirus (Accession: GQ927711) covering 87.23% of the query and demonstrating 74.10% pairwise identity to OP287812. Additionally, one BLASTn hit indicated homology to a partial Ghanaian *Eidolon helvum* kobuvirus sequence, which resolved as basal to canid and human Aichivirus A in phylogenetic analysis (Accession: JX885611, Peptide: L Peptide, Contig Length: 1,120bp, Query Coverage: 100%, Pairwise Identity: 96.60%)^49^. Most hits indicated homology to Aichivirus A variants and coincided with findings from our BLASTx search (**Supplemental Table 2**).

**Table 2.** Top BLASTn Hit for OP287812. Identity percentages, E-Values, corresponding hit lengths, and NCBI Accessions for the highest-ranking BLASTn hit between OP287812 and NCBI kobuvirus sequences. Nucleotide lengths for the whole OP287812 genome, including the ORF, individual proteins, and the UTRs, along with the predicted protease cleavage sites (in single-letter amino acid code) marking the start and end positions of each protein, are included. There were not hits observed for the 3’UTR.

| **Genome**<br>**Region** | **Start**<br>**(nt)** | **End**<br>**(nt)** | **Predicted**<br>**Protease**<br>**Cleavage**<br>**Site** | **Pairwise**<br>**Identity**<br>**%** | **E-**<br>**Value** | **NCBI**<br>**Accession** | **Hit**<br>**Length**<br>**(bp)** |
| --- | --- | --- | --- | --- | --- | --- | --- |
| Whole<br>Genome | 1 | 8263 | - | 74.10% | 0 | GQ927711 | 7240 |
| 5'UTR | 1 | 679 | - | 70.70% | 8.88E-60 | MK671314 | 657 |
| ORF | 680 | 7984 | - | 74.70% | 0 | MW292482 | 7016 |
| L | 680 | 1234 | Q/G | 96.60% | 0 | JX885611 | 554 |
| VP0 | 1235 | 2347 | Q/H | 77.10% | 0 | MN648601 | 1117 |
| VP3 | 2348 | 3016 | Q/A | 78.60% | 2.50E-155 | MK201778 | 668 |
| VP1 | 3017 | 3742 | Q/G | 72.20% | 5.28E-101 | KJ950958 | 724 |
| 2A | 3743 | 4150 | Q/G | 74.80% | 2.85E-63 | KJ950958 | 391 |
| 2B | 4151 | 4645 | Q/G | 76.10% | 8.83E-90 | MF175074 | 478 |
| 2C | 4646 | 5650 | Q/G | 78.10% | 0 | MG200054 | 995 |
| 3A | 5651 | 5929 | Q/A | 68.40% | 1.86E-08 | KJ934637 | 209 |
| 3B | 5930 | 6007 | Q/G | 78.30% | 3.94E-06 | MT610361 | 80 |
| 3C | 6008 | 6577 | Q/S | 76.00% | 8.40E-110 | GQ927706 | 569 |
| 3D | 6578 | 7984 | - | 81.10% | 0 | MH052678 | 1406 |
| 3'UTR | 7985 | 8263 | - | - | - | - | - |

### Phylogenetic analysis suggests common ancestry between bat kobuvirus OP287812 and Aichivirus A genotypes

We generated a nucleotide-based ML phylogeny for the *Picornaviridae* family, incorporating our newly identified *E. dupreanum* sequence (OP287812) alongside Picornavirus sequences obtained from NCBI (**Fig. 2)**. OP287812 clustered within the previously described kobuviruses, consistent with earlier BLAST analyses. Specifically, OP287812 localized within the Aichivirus A subclade of kobuvirus. This clade received strong support, with a bootstrap value of 100/10. Notably, previously described bat kobuvirus sequences (Accessions: KJ641691 and KJ641686)^18,50,51^ did not cluster with our Malagasy bat kobuvirus but instead grouped sister to the clade containing our sequence, with other species in the Aichivirus F clade. We also constructed a second ML phylogeny focusing specifically on sequences within the kobuvirus genus. This tree included previously identified bat kobuvirus sequences visualized in our Picornaviridae tree, along with additional bat kobuvirus sequences from Vietnam^11^ (**Supplemental Figure 1**). These sequences also clustered within the Aichivirus A subclade but grouped separately and were more derived compared to our *E. dupreanum* sequence. Alignment statistics between our novel Malagasy sequences and these previously identified bat kobuviruses can be found in **Supplemental Table 3**.

**Figure 2.**
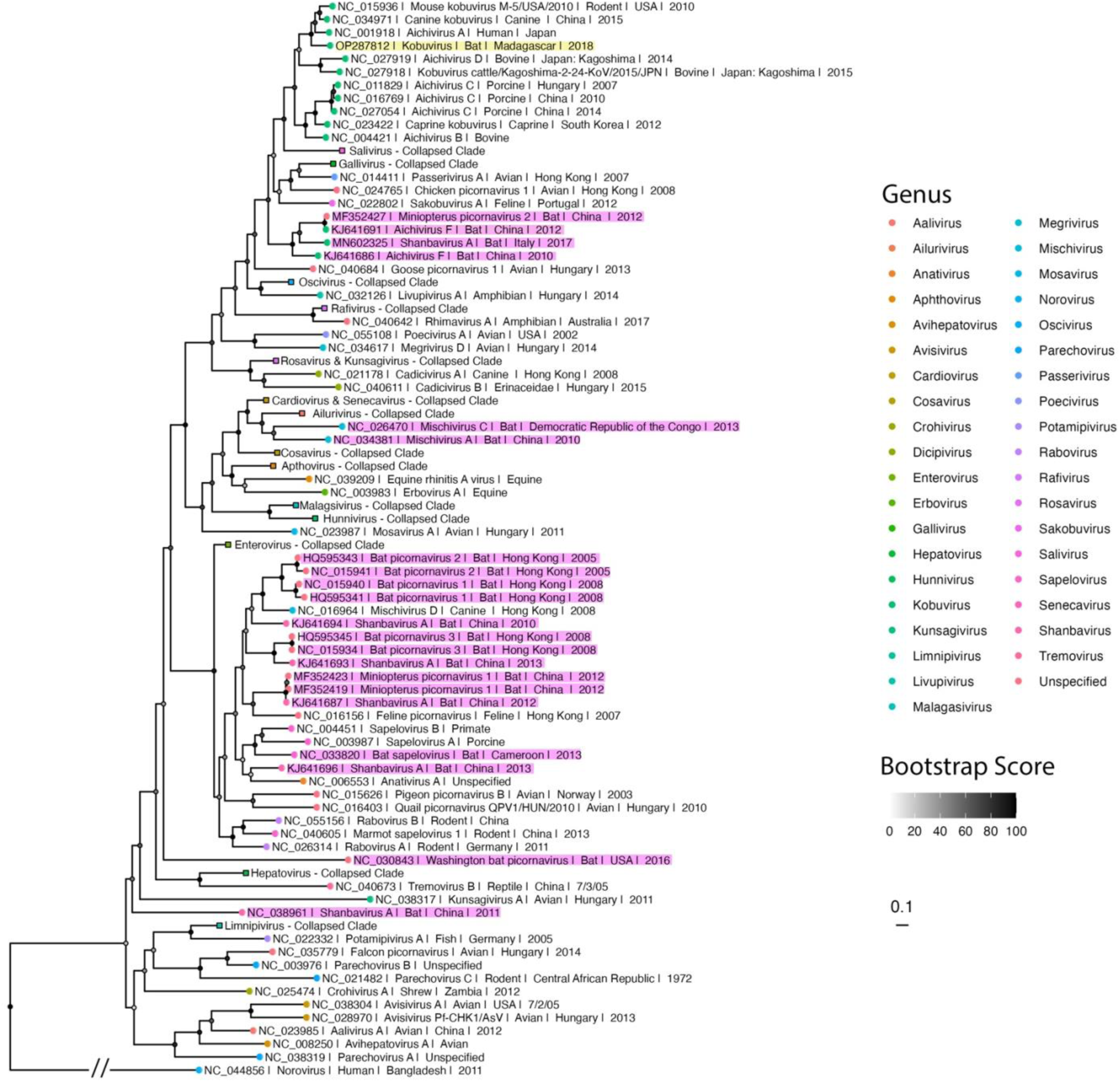
Phylogenetic analysis of OP287812 among previously identified picornaviruses. Maximum likelihood phylogeny of *Picornaviridae* sequences (nucleotide substitution model: TVM+I+G4). Node color, represented in greyscale, indicates bootstrap support, with darker shades corresponding to higher support values and lighter shades to lower support values. OP287812 is highlighted in yellow, while bat *Picornaviridae*, excluding OP287812, are highlighted in pink. Tip points are colored by broad viral taxonomic groups. Tip labels include NCBI accession number, virus species or genus, host, geographic origin, and year of identification, as available from NCBI. Branch lengths are scaled by nucleotide substitutions per site, noted by the scalebar. The tree is rooted with human norovirus (NC_044856). The branch length of this outgroup was shortened to improve phylogenetic tree visualization and is denoted as such with a double hash.

Our time tree (**Fig. 3**) estimated the most recent common ancestor (MRCA) for all kobuviruses to be in the year 1396 (∼628 years ago; 95% HPD: 1228–1556). It also supported the clustering of our Malagasy kobuvirus among Aichivirus A variants, with OP287812 diverging from its closest relatives at an MRCA dated to 1882 (∼142 years ago; 95% HPD: 1846–1914). This divergence occurred after an ancestral avian variant (GenBank Accession: KJ934637) diverged from the lineage approximately in 1857 (∼25 years prior; 95% HPD: 1814–1896). These divergence estimates suggest a relatively recent evolutionary history for kobuviruses

**Figure 3.**
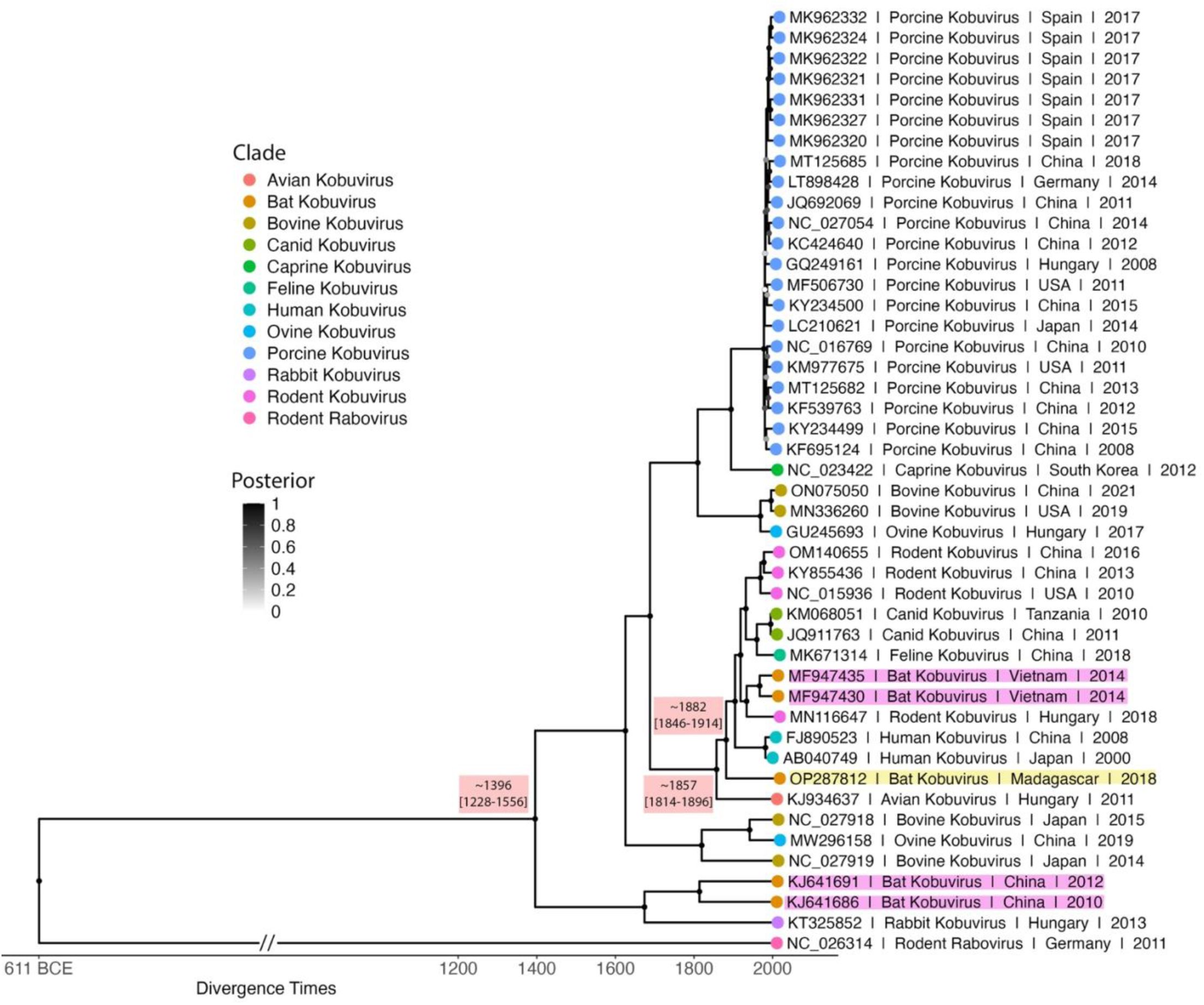
Bayesian phylogeny estimating time to MRCA for our novel *Eidolon dupreanum* Kobuvirus (OP287812) and other kobuvirus sequences. The Bayesian tree was constructed with 700 million runs of a strict molecular clock Bayesian Skyline Coalescent model (GTR+I+G4), implemented in BEAST2^45,52^. Node color, represented in greyscale, indicates posterior support, with darker shades corresponding to higher support values and lighter shades to lower support values after averaging of all 700 million trees after 10% burn-in. OP287812 is highlighted in yellow, while other bat kobuviruses are highlighted in pink. Tip points are colored by kobuvirus clade. Tip labels include NCBI accession number, clade, geographic origin, and year of identification, as available from NCBI. Estimated divergence times, with 95% highest posterior density (HPD) intervals, are depicted alongside key nodes. The outgroup is rodent rabovirus (NC_026314). The branch length of this outgroup was shortened to improve phylogenetic tree visualization and is denoted as such with a double hash.

## Discussion

Characterizing virus diversity in wildlife hosts serves as a foundational step towards downstream comprehensive analysis of host-virus ecology, transmission dynamics, and zoonotic risk. Bats are critical targets for pathogen surveillance due to their role as reservoirs for numerous zoonotic viruses, including coronaviruses, filoviruses, and henipaviruses. Effective surveillance in bat populations not only aids in detecting novel pathogens before they spill over into human populations but also provides key insights into the ecological and evolutionary factors that influence virus maintenance and transmission within and between species.

High genetic similarity between kobuviruses found in a variety of diverse mammalian hosts suggests the potential for interspecies transmission, with specific cross-species transmission events likely going undetected. For example, phylogenetic analyses provide evidence of shared kobuvirus ancestry between bat and rodent hosts^11^, bat and rabbit hosts^11^, as well as between cow and pig hosts^53^. More recently, compelling evidence for cross-species kobuvirus exchange between hosts has been identified on farms where a novel kobuvirus sequenced from sheep sampled in a paddock adjacent to a cattle farm, nested with viruses previously identified from the bovine clade^53,54^. Given the increasing evidence of kobuvirus circulation across diverse animal hosts, continuous surveillance in bat populations is essential for identifying potential spillover events early. Bat proximity to humans and livestock increases the opportunities for viral spillover^29^, and while kobuviruses have not yet been linked to major disease outbreaks, their evolutionary relationships with viruses in domestic animals suggest they could pose an emerging zoonotic threat. Understanding the prevalence, diversity, and transmission pathways of kobuviruses in bats could help mitigate this risk, providing critical insight into the dynamics of virus evolution and interspecies transmission.

Madagascar presents a unique environment for virus diversification due to its long isolation and dual African and Asian phylogeographic history. The high endemicity and unique diversity of Madagascar’s mammalian fauna, combined with widely-practiced wild meat consumption, rapid population growth, and multiple interacting anthropogenic threats, present considerable opportunity for viral crossover among diverse hosts^55–61^. We observe evolutionary relationships between Malagasy bat kobuviruses and Aichivirus A genotypes, including previously described bat kobuviruses^11^. Notably, our newly identified Malagasy bat kobuvirus appears basal to the vast mammalian host radiation^62^ that has since taken place within the Aichivirus A subclade. Our studies highlight the growing complexity of kobuvirus classification, suggesting that bat kobuviruses form a polyphyletic group across Aichivirus A and Aichivirus F clades^11,51^. This is especially concerning considering the potential for bat host coinfection with kobuvirus and other viruses with significant zoonotic potential, such as coronaviruses and henipaviruses^25,26^. Prior work has provided robust evidence for interspecies transmission of Aichivirus E kobuviruses from bats to rabbits^11^ and more muted evidence of cross species transmission of Aichivirus A kobuviruses from bats to rodents^11^. Broadly, high genetic similarity between kobuviruses found in a variety of mammalian hosts, including humans, suggests the potential for interspecies transmission, with specific cross-species transmission events likely going undetected. For example, phylogenetic analysis provide evidence of shared kobuvirus ancestry between bovine and porcine hosts, though definitive spillover has not yet been demonstrated^53^. Our findings fill a critical gap in understanding the diversity of kobuviruses in bats, emphasizing the need for enhanced surveillance in this unique ecological context.

Understanding the geographical distribution of bat kobuviruses is crucial for elucidating their evolutionary dynamics and potential zoonotic risks. A significant geographical bias in sampling is evident in the study of bat kobuviruses. For instance, previously described bat kobuvirus sequences used in our analyses (MF947429-MF947440 – *Scotophilus kuhlii*, KJ641691 – *Miniopterus fulginosus,* KJ641686 – *Myotis ricketti*) were exclusively recovered from fecal samples of Asian bat species^11,63^. Madagascar’s unique dual African and Asian evolutionary history may hold keys to understanding ancestral forms of bat kobuviruses and *Picornaviridae* more broadly. To fill these gaps, ongoing surveillance and additional sequencing data from diverse regions are essential. Such efforts could yield critical insights into the complex evolutionary histories and spatial dynamics that have shaped the trajectory of bat kobuviruses.

Our comparative analyses indicate conservation within genomic regions responsible for viral entry and replication^51,64,65^, such as VP1 and RdRP regions, between OP287812 and previously identified kobuviruses. Given these findings, it is crucial to explore the zoonotic potential of these Malagasy bat kobuviruses further. To advance our understanding, future studies should implement existing PCR protocols targeting the RdRp gene^64,66^ in RNA extracted from bat fecal samples. This approach will enable us to conduct longitudinal studies that build time series of infections, thereby elucidating the viral dynamics - such as transmission pathways and seasonal prevalence – that underlie the persistence of this pathogen in bat hosts.

## Conclusion

mNGS-based surveillance for bat viruses has resulted in incredibly diverse datasets, allowing for novel insights into wild bat virus ecology and evolution^25–27,63,67,68^. Here, we expand the known host and geographic range of kobuviruses to include *E. dupreanum* fruit bats of Madagascar, thereby expanding our understanding of bat-borne kobuviruses in the region. We describe the most complete bat kobuvirus genome identified to date, offering a glimpse into the origin and diversification of the kobuvirus genus more broadly. We find that bat kobuviruses in Madagascar phylogenetically nest among Aichivirus A genotypes and are highly divergent from previously described bat kobuviruses. Genome similarity analyses demonstrate significant conservation of kobuvirus genomic content across clades, particularly in regions of the virus genome involved in virus entry and replication. Further analyses are needed to determine whether this trend holds for other kobuviruses obtained from bats in various geographic landscapes, including unsampled species within Madagascar, or if the high identity between bat- and human-hosted kobuviruses identified here is unique to the region. While we did not identify kobuviruses in other sampled species (*P. rufus* and *R. madagascariensis*), these species should remain a focus of future sampling efforts, as their inclusion is essential for understanding the full ecological and evolutionary dynamics of bat kobuviruses. Moreover, our discovery of a partial kobuvirus genome in a second *E. dupreanum* bat highlights the need for more comprehensive sampling.

Given the high rates of human-bat contact in Madagascar^69^, our findings raise concern for public health. We strongly advocate for enhanced surveillance and detection efforts to further elucidate the ecology of these viruses in their wild bat hosts, as well as other animal hosts, as these efforts are critical for understanding their potential impact on both wildlife conservation and zoonotic disease transmission.

## Acknowledgements

We thank Anecia Gentles, Kimberly Rivera, Fifi Ravelomanantsoa, and Sarah Guth for help in the field and lab. We acknowledge the Virology Unit at the Institut Pasteur de Madagascar for logistical support, and we thank the Mention of Zoology and Animal Biodiversity at the University of Antananarivo and the Madagascar Ministry of the Environment and Sustainable Development for providing research and export permits. We thank Amy Kistler, Vida Ahyong, Angela Detweiler, Michelle Tan, and Norma Neff of the Chan Zuckerberg Biohub (CZB) for sequencing support and Cristina M. Tato, Maira Phelps, and Joseph L. DeRisi of CZB for logistical support. We thank the Brook lab at the University of Chicago for helpful contributions to the manuscript. This work was completed in part with resources provided by the University of Chicago’s Research Computing Center.

---

| Polyprotein | Start | End | Length | Predicted N-Terminal Cleavage Site | % Similarity to Reference Sequence | Reference Sequence Description |
| --- | --- | --- | --- | --- | --- | --- |
| 5'UTR | 1 | 679 | 679 |  |  |  |
| ORF | 680 | 7984 | 7305 |  | 71.37%; (JN387133) | Canine |
| L | 680 | 1234 | 555 | Q/G |  |  |
| VP0 | 1235 | 2347 | 1113 | Q/H | 71.17%; (FJ890523) | Aichivirus 1 |
| VP3 | 2348 | 3016 | 669 | Q/A | 78.42%; (FJ890523) | Aichivirus 1 |
| VP1 | 3017 | 3742 | 726 | Q/G | 67.72%; (JQ898342) | KobuV Sewage Isola |
| 2A | 3743 | 4150 | 408 | Q/G | 70.10%; (MH747478) | Canine |
| 2B | 4151 | 4645 | 495 | Q/G | 75.35%; (JQ898342) | KobuV Sewage Isola |
| 2C | 4646 | 5650 | 1005 | Q/G | 77.41%; (FJ890523) | Aichivirus 1 |
| 3A | 5651 | 5929 | 279 | Q/A | 65.37%; (MH747478) | Canine |
| 3B | 5930 | 6007 | 78 | Q/G | 67.07%; (JN387133) | Canine |

