## Supplementary Material for "Genomic characterization of novel bat kobuviruses in Madagascar: implications for viral evolution and zoonotic risk"

**Contents**

Supplemental Figure 1 – Kobuvirus Nucleotide ML Phylogeny 2

Supplemental Table 1 – Reference Sequences Used to Annotate OP287812 4

Supplemental Table 2 – Whole Genome BLASTx Hits for OP287812 5

Supplemental Table 3 – Genome alignment statistics of bat kobuviruses 7

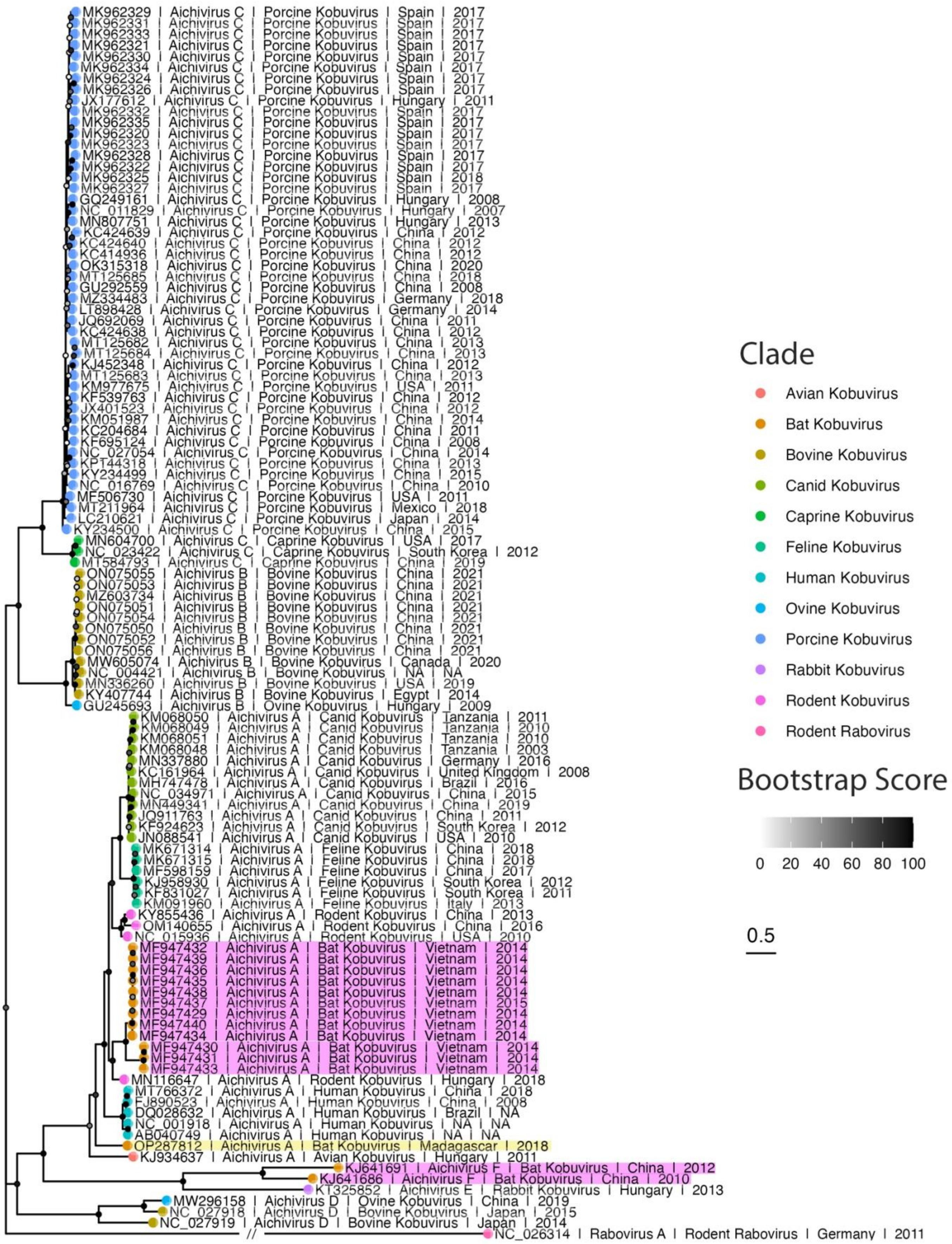
MK962329 | Aichivirus C | Porcine Kobuvirus | Spain | 2017
MK962331 | Aichivirus C | Porcine Kobuvirus | Spain | 2017
MK962333 | Aichivirus C | Porcine Kobuvirus | Spain | 2017
MK962321 | Aichivirus C | Porcine Kobuvirus | Spain | 2017
MK962330 | Aichivirus C | Porcine Kobuvirus | Spain | 2017
MK962334 | Aichivirus C | Porcine Kobuvirus | Spain | 2017
MK962324 | Aichivirus C | Porcine Kobuvirus | Spain | 2017
MK962326 | Aichivirus C | Porcine Kobuvirus | Spain | 2017
JX177612 | Aichivirus C | Porcine Kobuvirus | Hungary | 2011
MK962332 | Aichivirus C | Porcine Kobuvirus | Spain | 2017
MK962335 | Aichivirus C | Porcine Kobuvirus | Spain | 2017
MK962320 | Aichivirus C | Porcine Kobuvirus | Spain | 2017
MK962323 | Aichivirus C | Porcine Kobuvirus | Spain | 2017
MK962328 | Aichivirus C | Porcine Kobuvirus | Spain | 2017
MK962322 | Aichivirus C | Porcine Kobuvirus | Spain | 2017
MK962325 | Aichivirus C | Porcine Kobuvirus | Spain | 2018
MK962327 | Aichivirus C | Porcine Kobuvirus | Spain | 2017
GQ249161 | Aichivirus C | Porcine Kobuvirus | Hungary | 2008
NC_011829 | Aichivirus C | Porcine Kobuvirus | Hungary | 2007
MN807751 | Aichivirus C | Porcine Kobuvirus | Hungary | 2013
KC424639 | Aichivirus C | Porcine Kobuvirus | China | 2012
KC424640 | Aichivirus C | Porcine Kobuvirus | China | 2012
KC414936 | Aichivirus C | Porcine Kobuvirus | China | 2012
OK315318 | Aichivirus C | Porcine Kobuvirus | China | 2020
MT125685 | Aichivirus C | Porcine Kobuvirus | China | 2018
GU292559 | Aichivirus C | Porcine Kobuvirus | China | 2008
MZ334483 | Aichivirus C | Porcine Kobuvirus | Germany | 2018
LT898428 | Aichivirus C | Porcine Kobuvirus | Germany | 2014
JQ692069 | Aichivirus C | Porcine Kobuvirus | China | 2011
KC424638 | Aichivirus C | Porcine Kobuvirus | China | 2012
MT125682 | Aichivirus C | Porcine Kobuvirus | China | 2013
MT125684 | Aichivirus C | Porcine Kobuvirus | China | 2013
KJ452348 | Aichivirus C | Porcine Kobuvirus | China | 2012
MT125683 | Aichivirus C | Porcine Kobuvirus | China | 2013
KM977675 | Aichivirus C | Porcine Kobuvirus | USA | 2011
KF539763 | Aichivirus C | Porcine Kobuvirus | China | 2012
JX401523 | Aichivirus C | Porcine Kobuvirus | China | 2012
KM051987 | Aichivirus C | Porcine Kobuvirus | China | 2014
KC204684 | Aichivirus C | Porcine Kobuvirus | China | 2011
KF695124 | Aichivirus C | Porcine Kobuvirus | China | 2008
NC_027054 | Aichivirus C | Porcine Kobuvirus | China | 2014
KP144318 | Aichivirus C | Porcine Kobuvirus | China | 2013
KY234499 | Aichivirus C | Porcine Kobuvirus | China | 2015
NC_016769 | Aichivirus C | Porcine Kobuvirus | China | 2010
MF506730 | Aichivirus C | Porcine Kobuvirus | USA | 2011
MT211964 | Aichivirus C | Porcine Kobuvirus | Mexico | 2018
LC210621 | Aichivirus C | Porcine Kobuvirus | Japan | 2014
KY234500 | Aichivirus C | Porcine Kobuvirus | China | 2015
MN604700 | Aichivirus C | Caprine Kobuvirus | USA | 2017
NC_023422 | Aichivirus C | Caprine Kobuvirus | South Korea | 2012
MT584793 | Aichivirus C | Caprine Kobuvirus | China | 2019
ON075055 | Aichivirus B | Bovine Kobuvirus | China | 2021
ON075053 | Aichivirus B | Bovine Kobuvirus | China | 2021
MZ603734 | Aichivirus B | Bovine Kobuvirus | China | 2021
ON075051 | Aichivirus B | Bovine Kobuvirus | China | 2021
ON075054 | Aichivirus B | Bovine Kobuvirus | China | 2021
ON075050 | Aichivirus B | Bovine Kobuvirus | China | 2021
ON075052 | Aichivirus B | Bovine Kobuvirus | China | 2021
ON075056 | Aichivirus B | Bovine Kobuvirus | China | 2021
MW605074 | Aichivirus B | Bovine Kobuvirus | Canada | 2020
NC_004421 | Aichivirus B | Bovine Kobuvirus | NA | NA
MN336260 | Aichivirus B | Bovine Kobuvirus | USA | 2019
KY407744 | Aichivirus B | Bovine Kobuvirus | Egypt | 2014
GU245693 | Aichivirus B | Ovine Kobuvirus | Hungary | 2009
KM068050 | Aichivirus A | Canid Kobuvirus | Tanzania | 2011
KM068049 | Aichivirus A | Canid Kobuvirus | Tanzania | 2010
KM068051 | Aichivirus A | Canid Kobuvirus | Tanzania | 2010
KM068048 | Aichivirus A | Canid Kobuvirus | Tanzania | 2003
MN337880 | Aichivirus A | Canid Kobuvirus | Germany | 2016
KC161964 | Aichivirus A | Canid Kobuvirus | United Kingdom | 2008
MH747478 | Aichivirus A | Canid Kobuvirus | Brazil | 2016
NC_034971 | Aichivirus A | Canid Kobuvirus | China | 2015
MN449341 | Aichivirus A | Canid Kobuvirus | China | 2019
JQ911763 | Aichivirus A | Canid Kobuvirus | China | 2011
KF924623 | Aichivirus A | Canid Kobuvirus | South Korea | 2012
JN088541 | Aichivirus A | Canid Kobuvirus | USA | 2010
MK671314 | Aichivirus A | Feline Kobuvirus | China | 2018
MK671315 | Aichivirus A | Feline Kobuvirus | China | 2018
MF598159 | Aichivirus A | Feline Kobuvirus | China | 2017
KJ958930 | Aichivirus A | Feline Kobuvirus | South Korea | 2012
KF831027 | Aichivirus A | Feline Kobuvirus | South Korea | 2011
KM091960 | Aichivirus A | Feline Kobuvirus | Italy | 2013
KY855436 | Aichivirus A | Rodent Kobuvirus | China | 2013
OM140655 | Aichivirus A | Rodent Kobuvirus | China | 2016
NC_015936 | Aichivirus A | Rodent Kobuvirus | USA | 2010
MF947432 | Aichivirus A | Bat Kobuvirus | Vietnam | 2014
MF947439 | Aichivirus A | Bat Kobuvirus | Vietnam | 2014
MF947436 | Aichivirus A | Bat Kobuvirus | Vietnam | 2014
MF947435 | Aichivirus A | Bat Kobuvirus | Vietnam | 2014
MF947438 | Aichivirus A | Bat Kobuvirus | Vietnam | 2014
MF947437 | Aichivirus A | Bat Kobuvirus | Vietnam | 2015
MF947429 | Aichivirus A | Bat Kobuvirus | Vietnam | 2014
MF947440 | Aichivirus A | Bat Kobuvirus | Vietnam | 2014
MF947434 | Aichivirus A | Bat Kobuvirus | Vietnam | 2014
MF947430 | Aichivirus A | Bat Kobuvirus | Vietnam | 2014
MF947431 | Aichivirus A | Bat Kobuvirus | Vietnam | 2014
MF947433 | Aichivirus A | Bat Kobuvirus | Vietnam | 2014
MN116647 | Aichivirus A | Rodent Kobuvirus | Hungary | 2018
MT766372 | Aichivirus A | Human Kobuvirus | China | 2018
FJ890523 | Aichivirus A | Human Kobuvirus | China | 2008
DQ028632 | Aichivirus A | Human Kobuvirus | Brazil | NA
NC_001918 | Aichivirus A | Human Kobuvirus | NA | NA
AB040749 | Aichivirus A | Human Kobuvirus | NA | NA
OP287812 | Aichivirus A | Bat Kobuvirus | Madagascar | 2018
KJ934637 | Aichivirus A | Avian Kobuvirus | Hungary | 2011
KJ641691 | Aichivirus F | Bat Kobuvirus | China | 2012
KJ641686 | Aichivirus F | Bat Kobuvirus | China | 2010
KT325852 | Aichivirus E | Rabbit Kobuvirus | Hungary | 2013
MW296158 | Aichivirus D | Ovine Kobuvirus | China | 2019
NC_027918 | Aichivirus D | Bovine Kobuvirus | Japan | 2015
NC_027919 | Aichivirus D | Bovine Kobuvirus | Japan | 2014
//
NC_026314 | Rabovirus A | Rodent Rabovirus | Germany | 2011
Clade
Avian Kobuvirus
Bat Kobuvirus
Bovine Kobuvirus
Canid Kobuvirus
Caprine Kobuvirus
Feline Kobuvirus
Human Kobuvirus
Ovine Kobuvirus
Porcine Kobuvirus
Rabbit Kobuvirus
Rodent Kobuvirus
Rodent Rabovirus
Bootstrap Score
0 20 40 60 80 100
0.5

26

**Supplemental Figure 1 – Phylogenetic analysis of OP287812 among previously identified kobuviruses.** Maximum likelihood phylogeny of kobuvirus sequences (nucleotide substitution model: GTR+I+G4). Node color, represented in greyscale, indicates bootstrap support, with darker shades corresponding to higher support values and lighter shades to lower support values. Madagascar bat kobuvirus sequence OP287812 is highlighted in yellow, while other bat kobuviruses are highlighted in pink. Tip points are colored by kobuvirus clade. Tip labels include NCBI accession number, virus species, clade, geographic origin, and year of identification, as available from NCBI. Branch lengths are scaled by nucleotide substitution per site, noted by the scalebar. The tree is rooted with rodent rabovirus (NC_026314). The branch length of this outgroup was shortened to improve phylogenetic tree visualization and is denoted as such with a double hash.

| Genome Region | Start (nt) | End (nt) | % Similarity To NCBI Reference | NCBI Accession |
|---|---|---|---|---|
| 5’UTR | 1 | 679 | - | - |
| ORF | 680 | 7984 | 71.37% | JN387133 |
| L | 680 | 1234 | - | - |
| VP0 | 1235 | 2347 | 71.17% | FJ890523 |
| VP3 | 2348 | 3016 | 78.42% | FJ890523 |
| VP1 | 3017 | 3742 | 67.72% | JQ898342 |
| 2A | 3743 | 4150 | 70.10% | MH747478 |
| 2B | 4151 | 4645 | 75.35% | JQ898342 |
| 2C | 4646 | 5650 | 77.41% | FJ890523 |
| 3A | 5651 | 5929 | 65.37% | MH747478 |
| 3B | 5930 | 6007 | 67.07% | JN387133 |
| 3C | 6008 | 6577 | 75.74% | MH747478 |
| 3D | 6578 | 7984 | 80.98% | JN387133 |
| 3’UTR | 7985 | 8263 | - | - |

**Supplemental Table 1 – Reference Sequences Used for Annotating the OP287812 Genome.** Summary of reference sequences used to annotate the OP287812 genome and their corresponding NCBI accession number, including the percent similarity between the reference sequence and OP287812. Columns without data indicate regions that were manually annotated (i.e. 5’ UTR, L Peptide, 3’ UTR) without the use of a reference sequence.

| Genome Region | Start (aa) | End (aa) | Predicted Protease Cleavage Site | Pairwise Identity % | E-Value | NCBI Accession | Hit Length (bp) |
|---|---|---|---|---|---|---|---|
| Whole Genome | 1 | 2754 | - | 78.10% | 0 | AWK02691 | 2409 |
| ORF | 227 | 2661 | - | 78.10% | 0 | AWK02691 | 2409 |
| L | 227 | 411 | Q/G | 98.90% | 5.38E-128 | AGL97808 | 184 |
| VP0 | 412 | 782 | Q/H | 82.00% | 3.89E-164 | QUJ18037 | 372 |
| VP3 | 783 | 1005 | Q/A | 87.50% | 2.65E-119 | AVH76467 | 223 |
| VP1 | 1006 | 1247 | Q/G | 77.60% | 1.65E-106 | AVH76458 | 222 |
| 2A | 1248 | 1383 | Q/G | 74.30% | 6.06E-42 | QIE07158 | 135 |
| 2B | 1384 | 1548 | Q/G | 84.80% | 6.22E-87 | AVH76472 | 164 |
| 2C | 1549 | 1883 | Q/G | 86.90% | 0 | AFV70595 | 334 |
| 3A | 1884 | 1976 | Q/A | 64.90% | 1.23E-31 | QUJ18037 | 92 |

| | | | | | | | |
|---|---|---|---|---|---|---|---|
| 3B | 1977 | 2002 | Q/G | 88.90% | 1.53E-04 | AXE75324 | 25 |
| 3C | 2003 | 2192 | Q/S | 78.90% | 3.26E-101 | QEV86991 | 189 |
| 3D | 2193 | 2661 | - | 88.50% | 0 | YP_004782207 | 467 |

**Supplemental Table 2 – Top BLASTx Hit for OP287812.** Identity percentages, E-Values, corresponding hit lengths, and NCBI Accessions for the highest-ranking BLASTx hit between OP287812 and NCBI kobuvirus sequences. Amino acid lengths for the OP287812 genome, as well as predicted protease cleavage sites (in single-letter amino acid code) indicating start and end positions for individual proteins, are listed. No hits were observed within the 5’UTR and 3’UTR.

| **Sequence ID** | **Country** | **Host** | **Percent Identity (%)** | **Percent Coverage (%)** | **Mismatches** | **Length (bp)** |
|---|---|---|---|---|---|---|
| OR082796 | Madagascar | *E. dupreanum* | 96.39 | 24.53 | 52 | 2077 |
| MF947440 | Vietnam | *S. kuhlii* | 71.97 | 76.21 | 1724 | 6354 |
| MF947439 | Vietnam | *S. kuhlii* | 71.88 | 76.21 | 1730 | 6354 |
| MF947438 | Vietnam | *S. kuhlii* | 72.05 | 82.16 | 1843 | 6859 |
| MF947437 | Vietnam | *S. kuhlii* | 72.06 | 82.16 | 1842 | 6859 |
| MF947436 | Vietnam | *S. kuhlii* | 71.59 | 71.09 | 1628 | 5931 |
| MF947435 | Vietnam | *S. kuhlii* | 71.86 | 75.70 | 1719 | 6312 |
| MF947434 | Vietnam | *S. kuhlii* | 71.99 | 75.70 | 1711 | 6312 |
| MF947433 | Vietnam | *S. kuhlii* | 69.13 | 64.83 | 1629 | 5393 |
| MF947432 | Vietnam | *S. kuhlii* | 71.83 | 76.16 | 1732 | 6350 |
| MF947431 | Vietnam | *S. kuhlii* | 69.74 | 68.15 | 1679 | 5667 |
| MF947430 | Vietnam | *S. kuhlii* | 69.83 | 67.57 | 1659 | 5619 |
| MF947429 | Vietnam | *S. kuhlii* | 72.08 | 76.04 | 1713 | 6340 |
| KJ641691 | China | *M. fuliginosus* | 43.83 | 83.71 | 3228 | 7279 |
| KJ641686 | China | *M. ricketti* | 47.34 | 87.69 | 3219 | 7555 |

**Supplemental Table 3 – Genome alignment statistics between novel Madagascar bat Kobuviruses and previously identified bat kobuviruses with OP287812 set as the reference**

**sequence.** Alignment statistics between bat kobuvirus sequences used in **Supplemental Figure 1**.
